## Supplementary figures and images for "Cryo-EM structures and binding of mouse and human ACE2 to SARS-CoV-2 variants of concern indicate that mutations enabling immune escape could expand host range"

### S1-S10 Fig

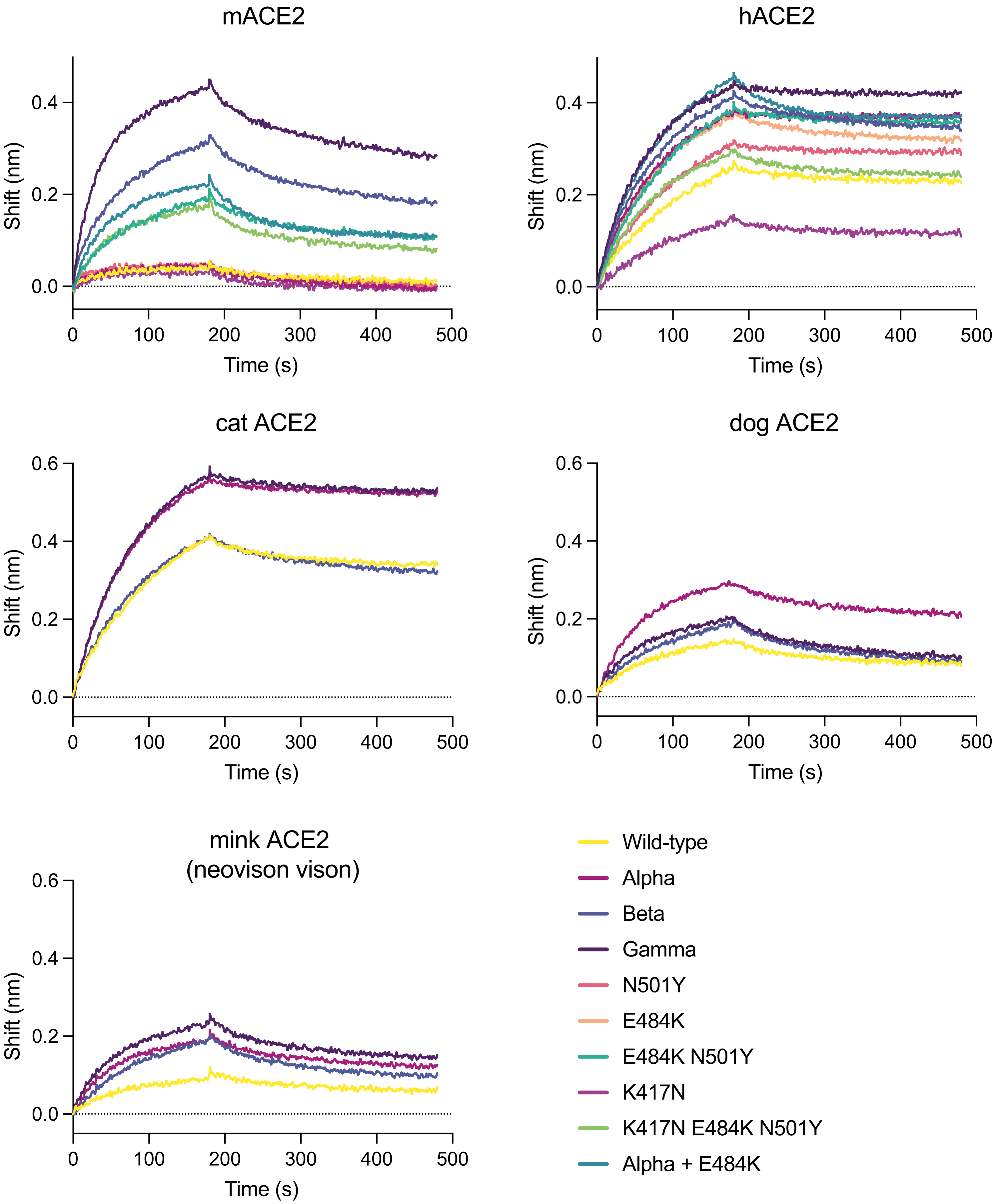

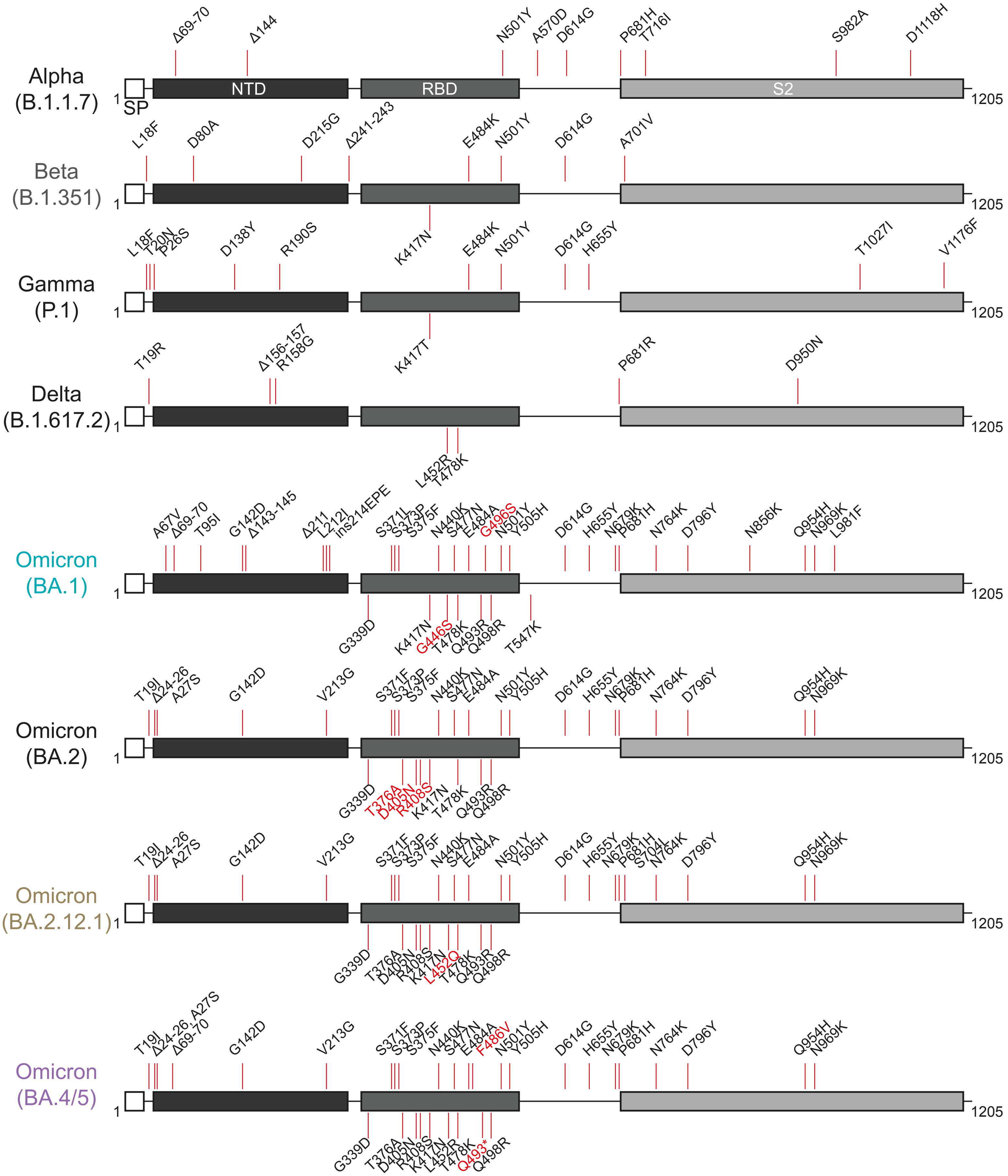

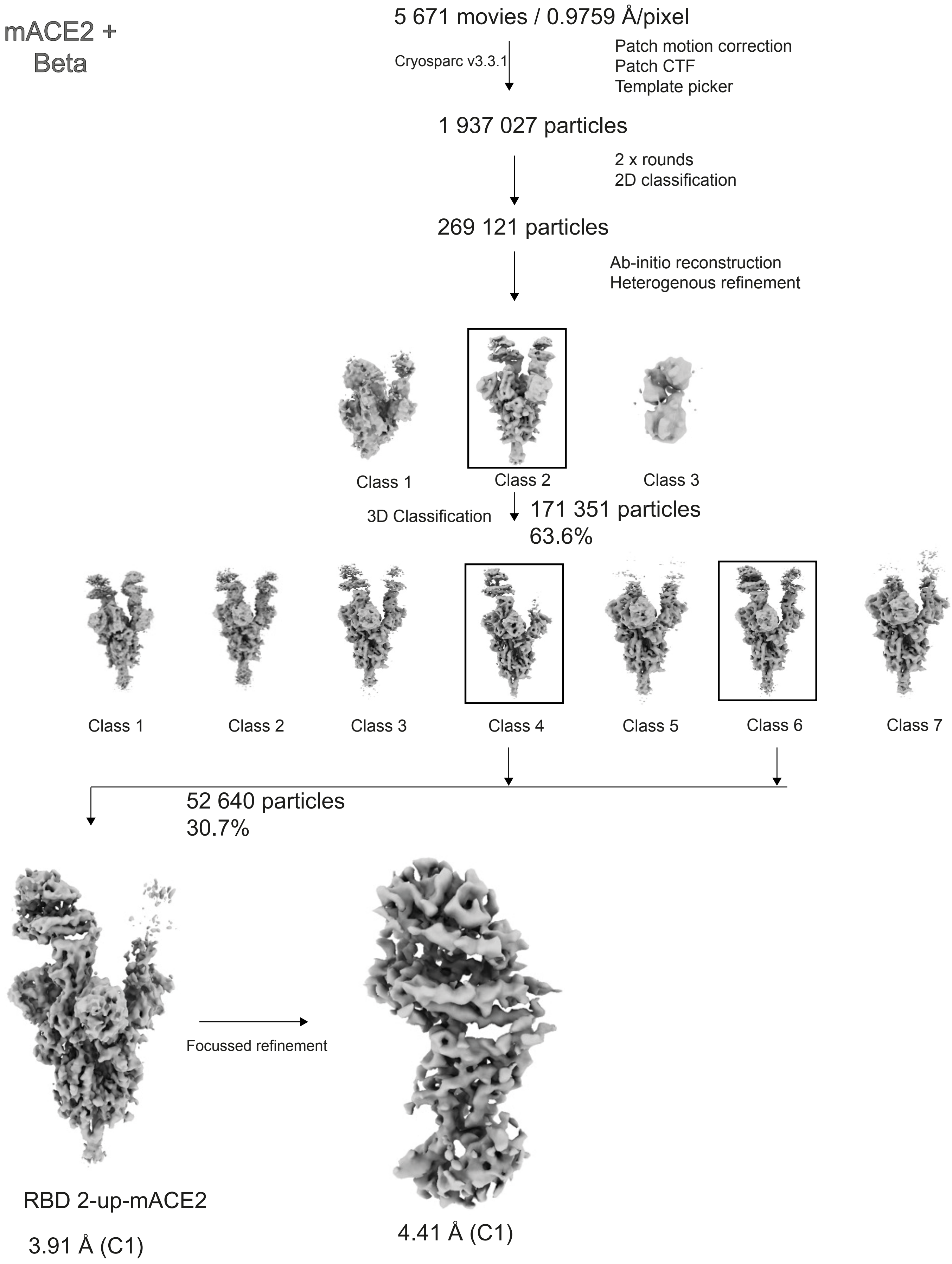

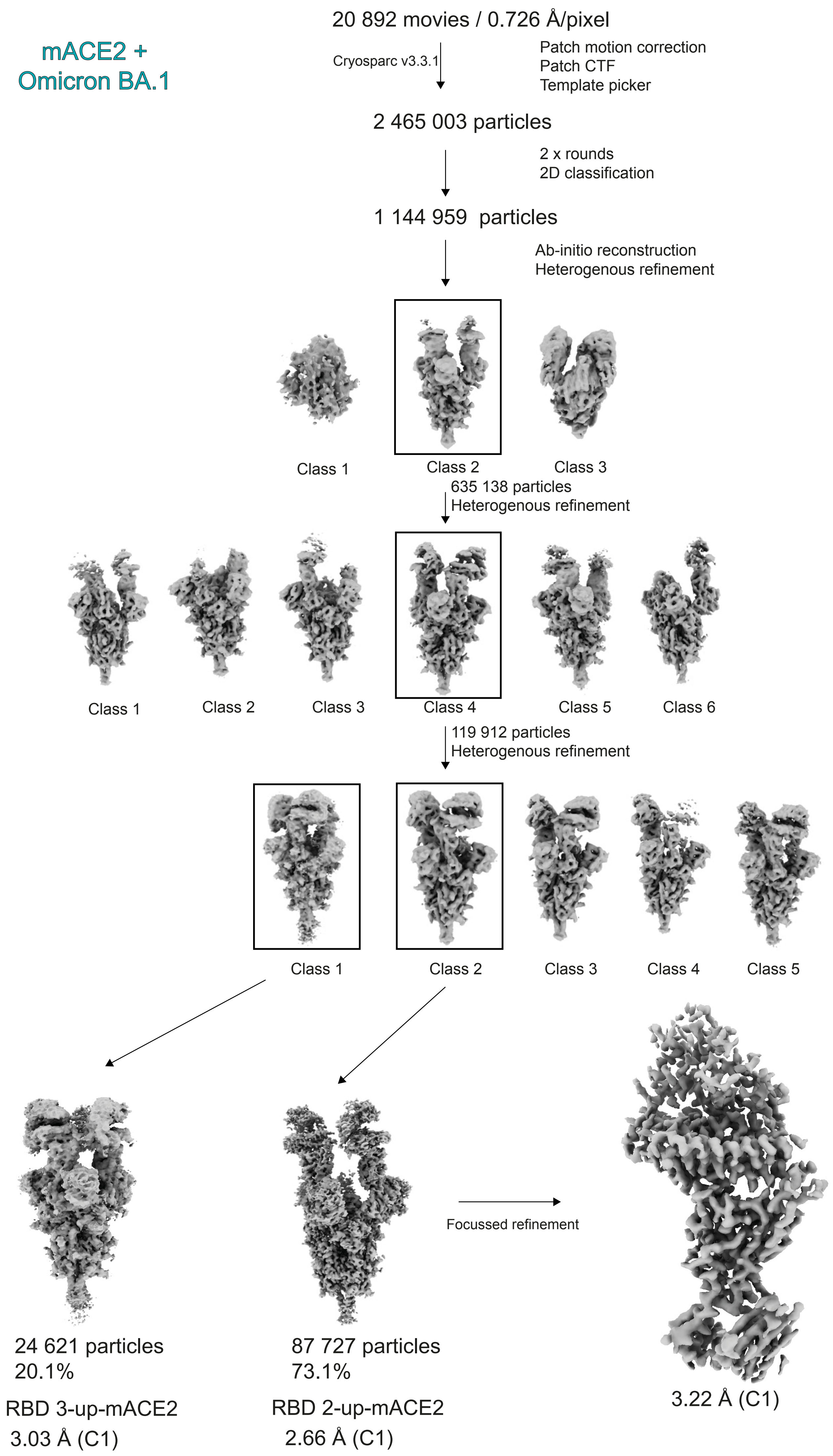

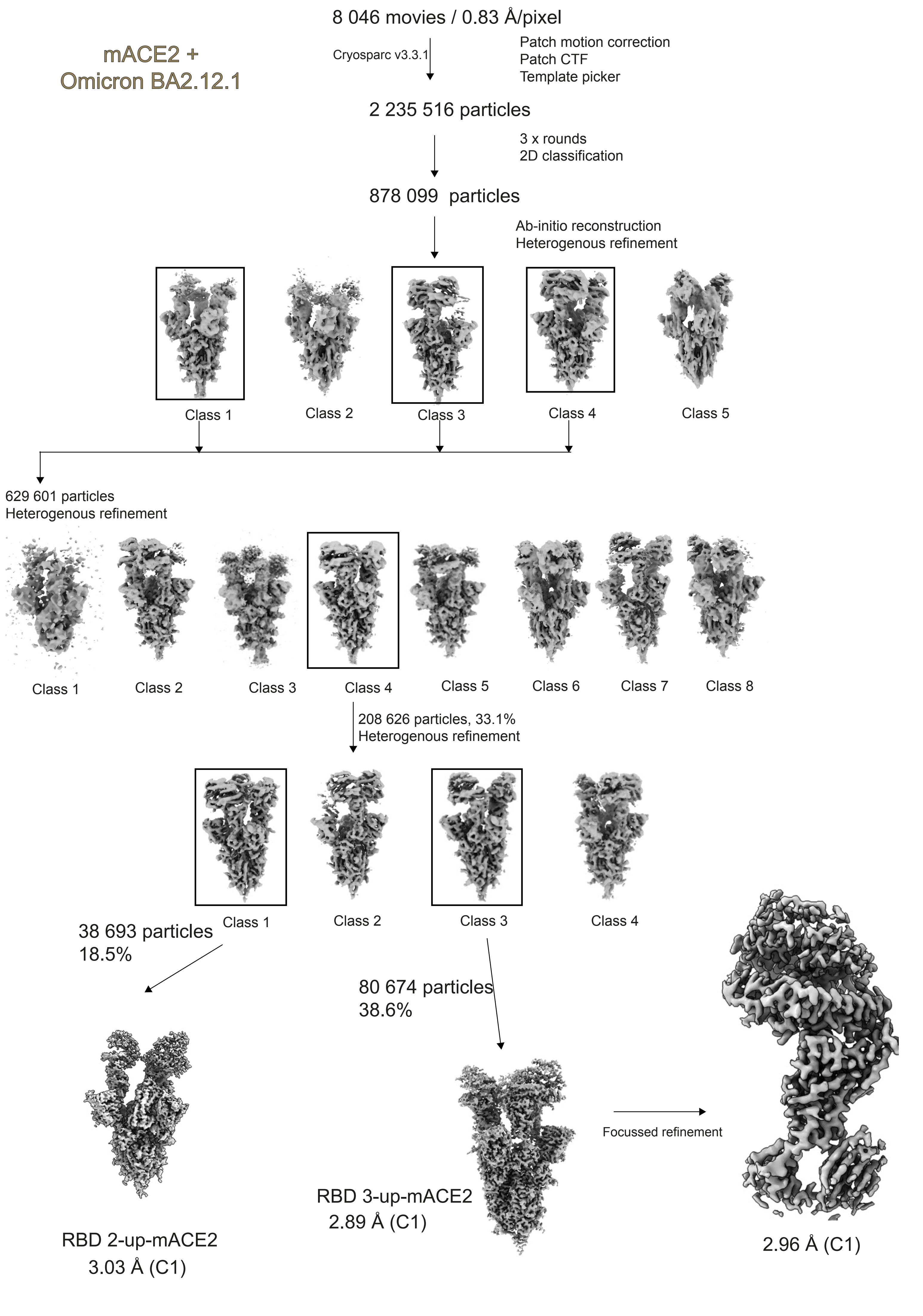

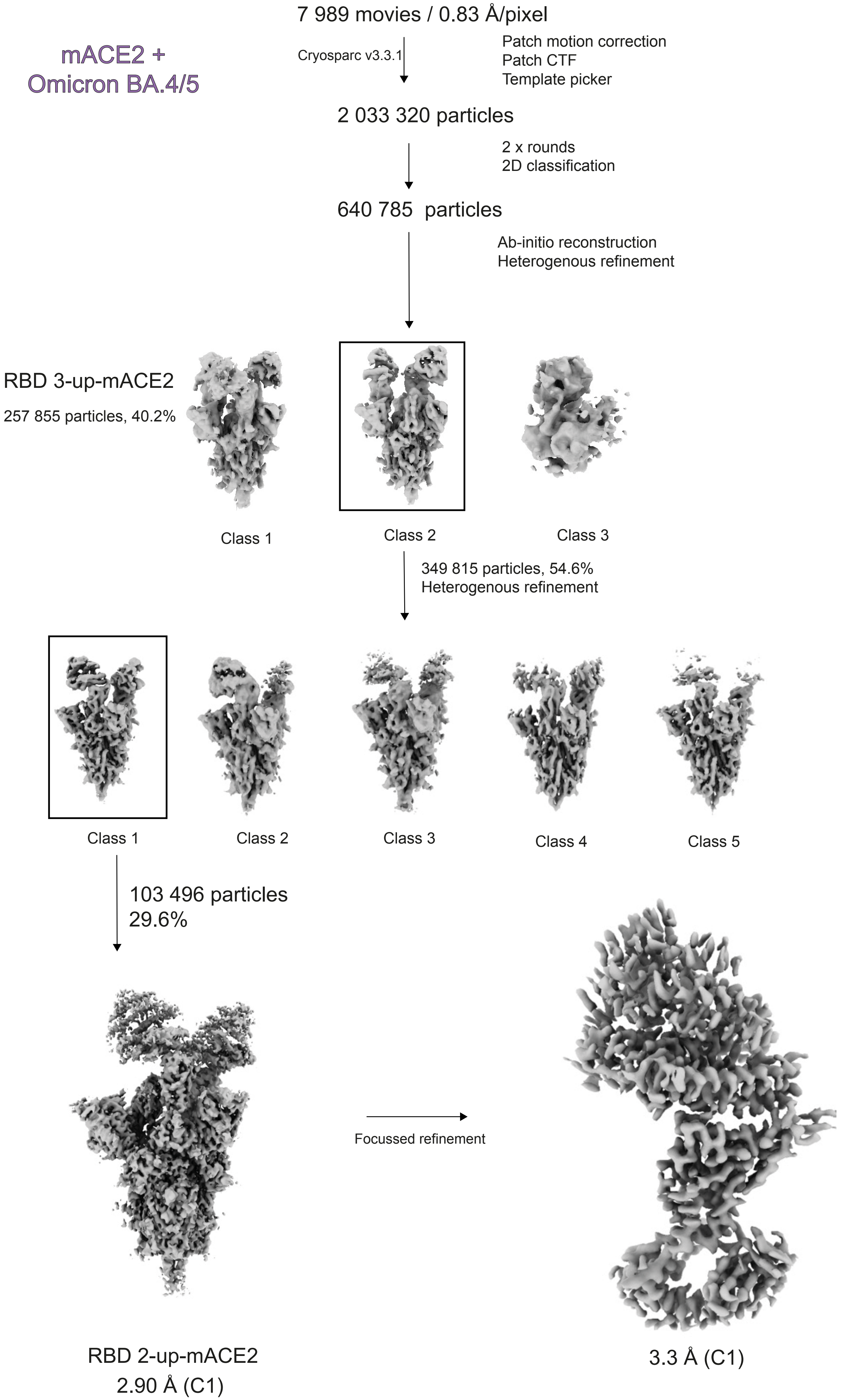

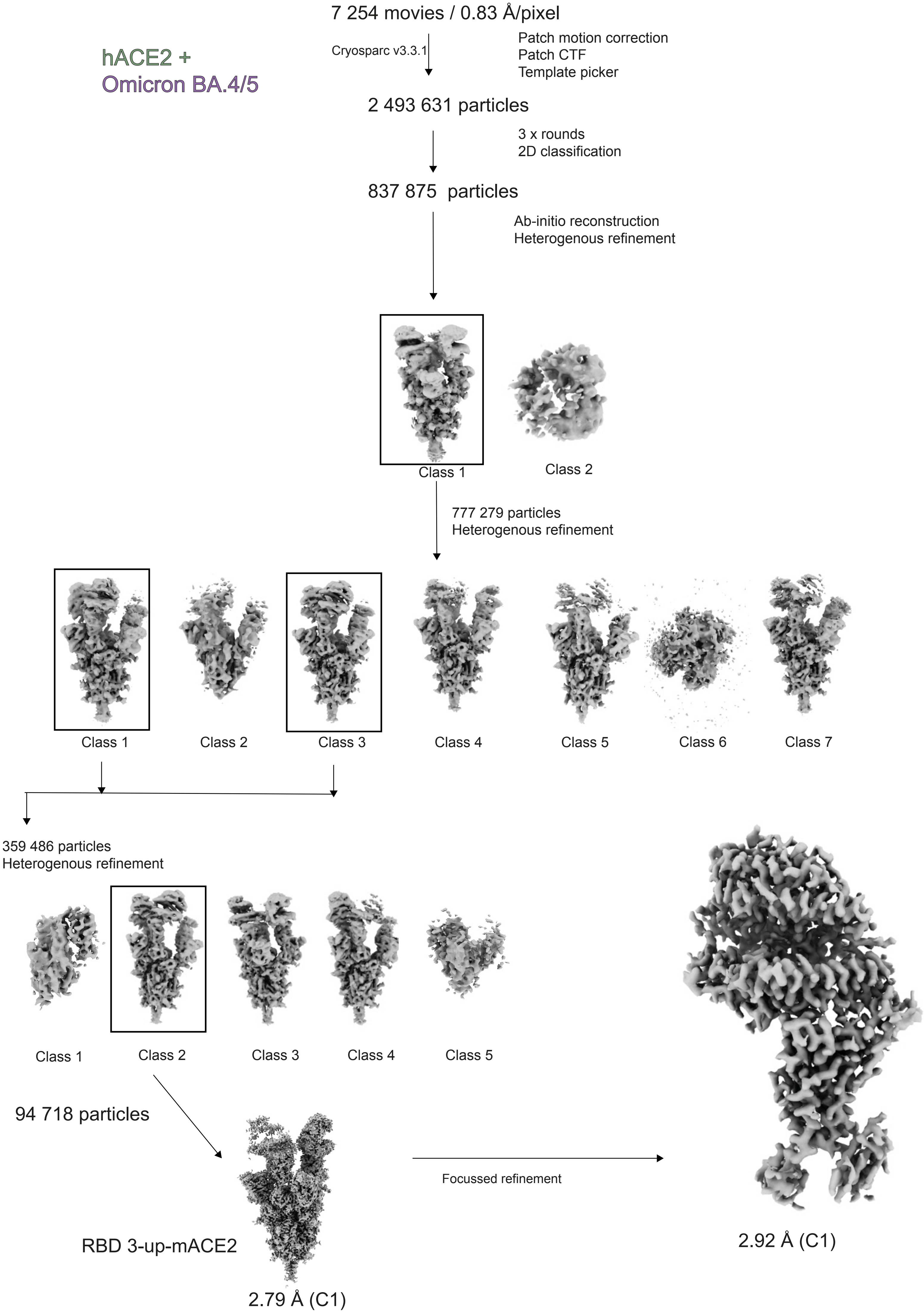

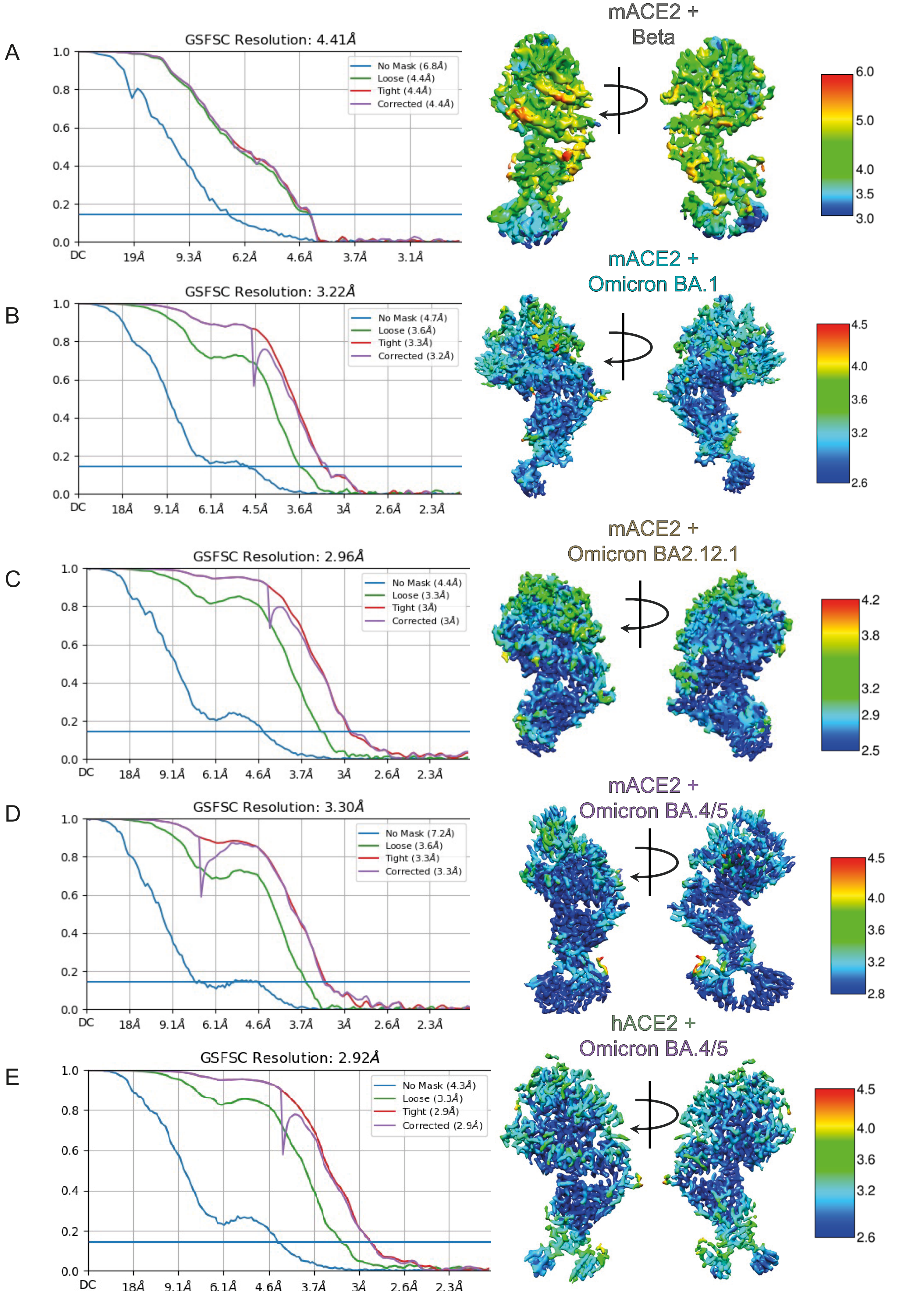

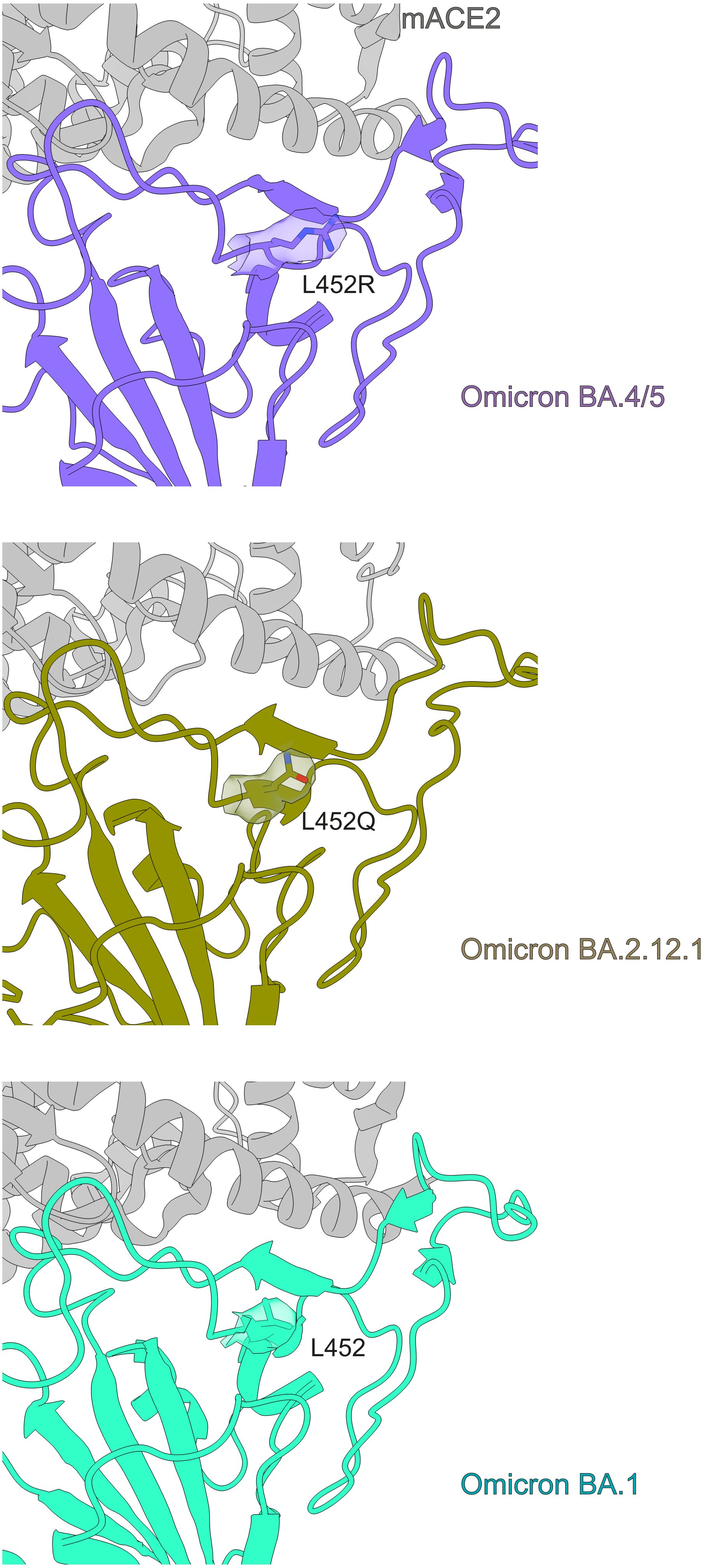

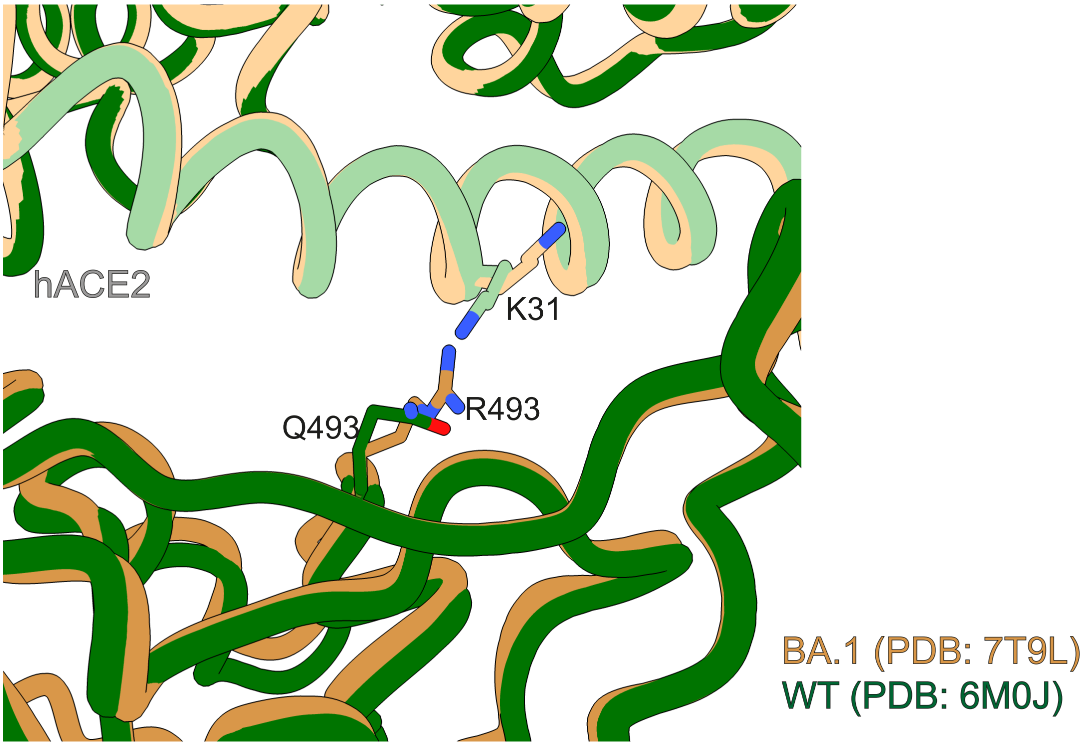
