## Supplementary material for "Cryo-EM structures and binding of mouse and human ACE2 to SARS-CoV-2 variants of concern indicate that mutations enabling immune escape could expand host range": S1 Table

| Data collection and processing | Omicron BA4/5-human ACE2 (EMDB-15588) (PDB 8AQS) | Beta-mouse ACE2 (EMDB-15589) (PDB 8AQT) | Omicron BA1-mouse ACE2 (EMDB-15590) (PDB 8AQU) | Omicron BA2.12.1-mouse ACE2 (EMDB-15591) (PDB 8AQV) | Omicron BA4/5-mouse ACE2 (EMDB-15592) (PDB 8AQW) |
|---|---|---|---|---|---|
| Magnification | 96kx | 150kx | 165kx | 96kx | 96kx |
| Voltage (kV) | 300 | 200 | 300 | 300 | 300 |
| Microscope | TFS Titan G4 | TFS Talos Arctica | TFS Titan G4 | TFS Titan G4 | TFS Titan G4 |
| Electron exposure (e–/Å$^2$) | 60 | 40 | 60 | 60 | 60 |
| Defocus range (-μm) | 0.8-2.5 | 0.8-2.5 | 0.7-2.0 | 0.7-2.4 | 0.8-2.4 |
| Pixel size (Å) | 0.83 | 0.9759 | 0.726 | 0.83 | 0.83 |
| Symmetry imposed | C1 | C1 | C1 | C1 | C1 |
| Initial particle images (no.) | 837 875 | 269 121 | 1 144 959 | 878 099 | 640 795 |
| Final particle images (no.) | 94 718 | 52 640 | 87 727 | 80 674 | 103 496 |
| Map resolution (Å) | 2.92 | 4.41 | 3.22 | 2.96 | 3.3 |
| FSC threshold | 0.143 | 0.143 | 0.143 | 0.143 | 0.143 |
| **Refinement** | | | | | |
| Initial model used (PDB code) | n/a | 7QO7, 7FDK | 7QO7, 7FDG | n/a | n/a |
| Map sharpening B factor (Å2) | -68.4 | -135.8 | -78.8 | -63.1 | -46.6 |
| Model composition | | | | | |
| Non-hydrogen atoms | 6493 | 6321 | 6340 | 6448 | 6458 |
| Protein residues | 788 | 777 | 777 | 789 | 789 |
| Water | 1 | 0 | 0 | 0 | 0 |
| Ligands | NAG:6 ZN:1 | NAG:3 | NAG:3 | NAG:3 | NAG:3 |

| | | | | | |
|---|---|---|---|---|---|
| *B* factors (Å$^2$) (lowest/highest/mean) | 21.76/204.34/80.80 | 70.88/351.28/158.61 | 11.478/220.77/92.89 | 17.0/194.06/83.54 | 27.46/164.34/75.76 |
| Protein | 82.05/134.60/103.02 | 134.44/259.41/206.11 | 65.22/118.96/92.25 | 54.59/130.61/88.56 | 71.27/145.17/101.80 |
| Ligands | 6.58/6.58/6.58 | --- | --- | --- | --- |
| Water | | | | | |
| R.m.s. deviations | | | | | |
| Bond lengths (Å) | 0.002 (0) | 0.003 (0) | 0.002 (0) | 0.003 (1) | 0.002 (0) |
| Bond angles (°) | 0.522 (1) | 0.652 (4) | 0.606 (5) | 0.701 (12) | 0.582 (3) |
| Validation | | | | | |
| MolProbity score | 1.39 | 1.87 | 1.95 | 1.61 | 1.76 |
| Clash score | 4.10 | 9.08 | 8.47 | 4.53 | 6.12 |
| Poor rotamers (%) | 0.00 | 0.15 | 0.15 | 0.00 | 0.15 |
| Ramachandran plot | | | | | |
| Favored (%) | 96.81 | 94.31 | 91.72 | 94.39 | 93.63 |
| Allowed (%) | 2.93 | 5.56 | 8.28 | 4.97 | 6.11 |
| Disallowed (%) | 0.26 | 0.13 | 0.00 | 0.64 | 0.25 |

**S1 Table: Cryo-EM data collection, refinement, and validation statistics**
